# Assessing the antimicrobial activity of *Aframomum melegueta and Piper guineense* extract on pathogens of rot diseases of cucumber fruit

**DOI:** 10.1101/2024.09.21.614230

**Authors:** Patience Saturday Akpan, Eyoanwan Offon Umoren, Ebere Ifiokobong William, Aniefon Alphonsus Ibuot

## Abstract

The inhibitory potential of *Aframomum melegueta and Piper guineense* extract against the pathogens of fruit rot diseases of cucumber was assessed. Two fungal isolates isolated from rotten cucumber fruits were; *Fusarium oxysporium* and *Penicillium* sp. The test plant extract were prepared in concentrations of 100% and 70%. The antimicrobial potential of ethanolic extract of *Piper guineense* at 100% concentration against *Fusarium oxysporium* revealed inhibition zone of 26.6 mm while 70% concentration of same plant extract had 17.0 mm. Ethanolic plant extract of *A. melegueta* at 100% concentration showed 7.5 mm and 70% concentration of same plant extract had 6.5 mm. Aqueous plant extract of both plants showed very minimal inhibitory effect on *F. oxysporium* which ranged from 0.00mm to 8.5mm. For *Penicillium* sp, ethanolic extract of *Piper guineense* at 100% showed inhibition of 18.0mm and at 70% concentration of same plant extract showed inhibition of 12.5mm. Ethanolic plant extract of *A. melegueta* at 100% concentration inhibited the growth of *Penicillium* sp with inhibition zone of 15.5mm and the 70% concentration of same plant extract showed 11.5mm inhibition zone. The aqueous plant extract at 100% revealed inhibition zone of 15.5 mm and 70% concentration showed no inhibition. The phytochemical screening of test plants revealed the presence of most of the phytochemicals tested in abundance with very few exception.

## Introduction

Cucumber (*Cucumis Sativus*) which bears cucumform fruits is a extensively -cultivated crawling vine plant that is in the family of cucumbitaceae gourd mostly used as vegetables. The three main types of cucumber that are presently available are -slicing, picking and burpless /seedless; several cultivars have been created from these [1,2]. Cucumber (Cucumis sativus) which belongs to the family of Cucurbitaceae is a majorly -cultivated creeping vine plant that bears fruits often in cylindrical shape, and are used as cooking vegetables. Cucumber obviously considered an annual plant, have three main types/varieties —seedless, slicing and pickling, many cultivars have been created within these varieties [2, 3]. Cucumber has its origin from India and several varieties have been produced being that the plant has been cultivated for at least 3,000 years. Cucumbers are observed to have a high water content, mild and refreshing taste. They are very pleasant to eat and can help reduce dehydration in hot weather. People eat cucumber as a delicious and tasty food. [4]. Some consume cucumbers raw or pickled. Mature raw cucumbers give relief to persons who are suffering from celiac disease, and enhance skin health. The seeds of cucumber contains edible oil which can be extracted and used for cooking.

Dysentery can be treated using cooked immature cucumber. Its ability to soften the skin makes cucumber fruit significant in the cosmetic industry. A moist mass made from fresh cucumbers can be bring a relief to burns and open sores when applied. Parasitic worms can be expelled using cucumber seeds. The juice from cucumber leaves has been reported to induce vomiting and aid digestion. [5].

Cucumber is very vulnerable to *Fusarium Oxysporum* which causes *cucumberinum*. The initial manifestation begins six to eight weeks followed by the presence of pale yellow lesions at the stem base. These lesions most times expand and spread to cause a root and stem rot which causes rot and colonization of stem by the fungus leads to the breakdown of cortical tissues as the disease progresses. This will negatively influence the fruit yield of the plants.

The rot of fruits starts from the stem and commence generally as the fruits ripens. The fungal pathogen invades the flesh of the fruits causing discoloration of the tissue, deterioration and at times derogatory odour thus reducing the quality of fruit. Fungal entries into the fruit are caused by other pathogens, insects that assist in penetration and most times wind. Fruits rot of cucumber can be initiated as a result of dry weather. Majority of this disease gets into the field through transplanting of already infected plant or infected seed. Post-harvest rot is initiated on cucumber during packaging, storage and sometimes transportation of the products. The main causes of cucumber loss are due to disease caused by pathogenic microorganisms [6]

During the growing phase, infection by microorganisms (fungi and bacteria) may occur. Cucumber can be exposed to microbial infections during harvest season, handling after the harvest marketing and after purchase by the customer. Some microorganisms can cause disease in a broad range of plant species that are unrelated, and others are limited to a single host. Most bacteria gains entry through injuries incurred during harvest or natural openings and then initiates the spoilage. While fungi are capable of penetrating directly into the intact cuticle, stem, leaves, and fruits. When unripe, many cucumber fruits are resistant to fungal attack, the process of infection is stopped almost as soon as it has started. The ripening process is followed by a weakening of the cell wall and a decrease in the ability to synthesis an anti-fungal substance meant to protect the fruit from fungal attack, until at the end the fruits is no longer able to resist the attack of the fungus. The fruit decay begins as a results of increased respiration of the host tissue and heat production. Some molds have been reported to produce ethylene by themselves and this in turn makes healthy tissues susceptible to infection. Environmental factors such as humidity, temperature and atmospheric components influence the interactions between the plant tissues and microorganisms [7]

Apart from fruit rot disease, fungi and bacteria also causes fruit spot. A fruit spot is a definite localized area infection. The spot mostly enlarges and combines together to form a rot, at the injury or stem end is commonly where softening and discoloration begins which often cause a disintegration of tissues and then the infection. The infecting microorganisms are usually transmitted by insects through the injury points. The development of spot and rot is often rapid in warm and moist storage conditions. *Cladosporium cucumerium* a fungus which causes fruit cap disease of cucumber reveals dark appearances of lesion appearing corky on the fruits.

Some plants are embedded with components that are inhibitory to pathogens when extracted and applied on infected crops, these formations are considered as botanical tools that can be relied on for the prevention of cucumber rot caused by microorganisms. A vast array of botanicals plant have limitless ability to metabolites, most of which are phenols or their oxygen substituted derivatives. Some of these components include quinones, phenols, phenolic acids, flavones, flavonoids and tanins [8]. These groups of plant compounds exhibit antimicrobial effect and serves as a defense mechanisms of plant against pathogenic microorganisms. Plant parts like crude sap, volatile and essential oil extracts from the whole plant or plant parts like flowers, leaves, stem, fruits, seeds and root are widely used in preparing the antimicrobial compounds which are mostly applied against the different plant pathogens. The use of plant extracts against pathogens triggers plants latent defense mechanism in responses to infection by pathogenic microorganisms and also an aqueous plant extract which is rich in both phytochemical compounds and micro-organisms. Therefore this plant extract application can protect plants from disease causing organisms and also protect plants from seed-borne pathogen [9]

## 2.2 METHODS

### 2.2.1 Sample collection

The rotten cucumber fruit was collected from otor market, Ikot Ekpene, Akwa Ibom State and was brought to the microbiology laboratory for analysis.

### 2.2.2 Medium preparation

The medium used was potato dextrose agar (PDA) and was prepared according to manufacturer’s instructions. After preparation, the flask containing the medium was corked tightly using aluminum foil. A piece of aluminum foil was then used to wrap round the mouth of the flask and was sterilized with an autoclave at 121°C for 15minutes and was allowed to cool at about 45°C.

### 2.2.3 Isolation of fungi causing cucumber fruit rot

The bench area to be used was sterilized using 70% ethanol, the rotten cucumber fruit was also sterilized using 70% ethanol especially the point where the rot was. A sterile knife was used to cut the rotten portion to be used and ground using laboratory mortar and pestle. After which, 1g of the ground cucumber was weighed out and a tenfold serial dilution was carried out from 10^-1^ to 10^-5^ dilution factors A calibrated pipette was used to pipette 1ml from 10^-3^ and 10^-5^ dilution factor into two sterile petri dishes respectively. About 15ml of already prepared medium was poured into each of the petri dishes and was swirled to homogenize and was allowed to gelatinize. Thereafter the plates were wrapped using aluminum foil and labeled properly and kept at room temperature for 5days. After 5days, the plates were observed.

### 2.2.4 Characterization and Identification of fungal isolates

The fungal isolates were observed and characterized based on microscopic and macroscopic appearance, and their identification were made according to Ahmed and Ravinder Reddy [10].

### 2.2.5 Preparation of subculture

Potato Dextrose Agar (PDA) was prepared and poured into petri dishes and allowed to gelatinize. Each of the colonies isolated were inoculated in the plate and kept at 27°C room temperature for 5days.

### 2.2.6 Preparation of seeds Extracts

Two different plant species were used as treatment

(i) Ashanti pepper------- *Piper guinense*

(ii) Alligator pepper------- *Aframomum melequeta*

The seeds of the plant materials free from disease were dried under the sun for few days. The plant seeds were ground separately into fine powder and stored in a sterile transparent polythene bag. For further use.

#### 2.2.6.1 Water extract

The ground *Piper guineense* was put into a sterile plate with cover 50ml (wt/vol) of distilled water was put into the plate containing the ground *Piper guineense*. It was soaked for 24 hours and was filtered using what man’s filter paper. The filtrate was evaporated to dryness in water bath at about 45°C and the residue was kept and labeled in sample bottle and refrigerated at 4°C for preservation. The same procedure was carried out on seeds of *Aframomum melegueta*.

#### 2.2.6.2 Ethanol extract

The ground *Piper guineense* was put into a sterile plate with cover, 50ml of 70% (wt/vol) ethanol was put into the plate containing the ground *Piper guineense*. It was soaked for 72hours. After 72hours, it was filtered using what man’s filter paper. The filtrate was evaporated to dryness in water bath at about 45°C and the residues was discarded the evaporated extract was stored in a well labeled sample bottle and refrigerated at 4°C for preservation. The same procedure was carried out on seeds of *Aframomum melegueta*.

### 2.2.7 Preparation of the Disc (whatman’s perforated paper) with extracts

The whatman’s filter paper was perforated aseptically and placed in a sterile bottle with cork and was sterilized using hot air oven for 60 min at 160 °C, it was allowed to cool for 60minutes and was ready for use.

Solution of extracts was prepared with three concentrations, 70%, 100% for all the seed extracts both ethanol and aqueous.

### 2.2.8 Effect of plant extracts on radial growth of the test pathogens

Antifungal test was carried out on the pathogens using both water and ethanol extracts. Molten potato dextrose agar (PDA) (45^0^ C) was poured into six sterile petri dishes. They were allowed to gelatinize. *Fusarium oxysporum* was inoculated heavily on the plates containing PDA by streak plate method. Immediately, control disc, ethanol and aqueous extract of *Piper guinense* (Ashanti pepper) placed on the inoculated plates using sterile forceps. The same procedure was carried out on *Penicillium* sp. These same procedures were carried out for *Aframomum melequeta (*Alligator pepper) extracts. All the plates were kept at 26°C to 28°C.

### 2.2.9 Pathogenicity test

Healthy fruits with neither wounds nor blemishes were carefully selected for the pathogenicity test. The fruits were surface sterilized by swabbing with 70% ethanol and then in changes of sterile distilled water. A sterile knife was used to cut the cucumber fruits aseptically and holes were cut out from the cucumber fruits by means of 3mm diameter sterile cork borer. Each of the test pathogens were inoculated into each of the cucumber separately, the cores of tissues cut were replaced and the wound sealed with candle wax. Each fruit was tied up in a sterile transparent polythene bags to prevent entry of other organisms and kept at one point only, at 27°C. Controls consist of sterile PDA introduced into similarly made holes or incisions on healthy fruit, tied and kept at 27°C. After the period of seven days, the fruits were cut across by means of a sterile knife at the point of inoculation and the rot symptoms noted, the microorganisms re-isolated on the PDA from the advancing edge of the lesion were considered pathogenic.

### 2.3 PHYTOCHEMICAL SCREENING OF *Piper guineense* AND *Aframomum melegueta*

Phytochemical screening consist of chemical test which detect the presence of the tannins, saponins, alkaloids, flavonoids and gardiac glycoside in plant. The screening was carried out using Trease and Evans [11], Sofowora [12] and Harbone [13] methods.

## 3.0 RESULT AND DISCUSSION

### 3.1 RESULT

#### 3.1.1 Characterization and identification of fungal isolates

The fungal isolates identified were *Penicillium* sp. and *Fusarium oxysporum*. Ahmed and Ravinder Reddy [10] culture method was used for microscopic study of the isolates which are by growth colony appearance, surface color, Reverse color, Hyphae/mycelium, conidia, conidial head, vesicle.

#### 3.1.2 Effect of plant extracts on radial growth of the test pathogens

The effect of *Piper guineense* (Ashanti pepper) and *Aframomum Melegueta* (Alligator pepper) extract on *Fusarium oxysporum* and *Penicilliun* sp. were carried out. (Table 2) shows that at 100% concentration *Aframonum Melegueta* ethanolic extracts had 7.5mm zone of inhibition or effect on *Fusarium oxysporum* and 15.5mm on *Penicillium* sp. *Aframomum melegueta* aqueous extract had no inhibition on both isolates.

**Fig. 1:**
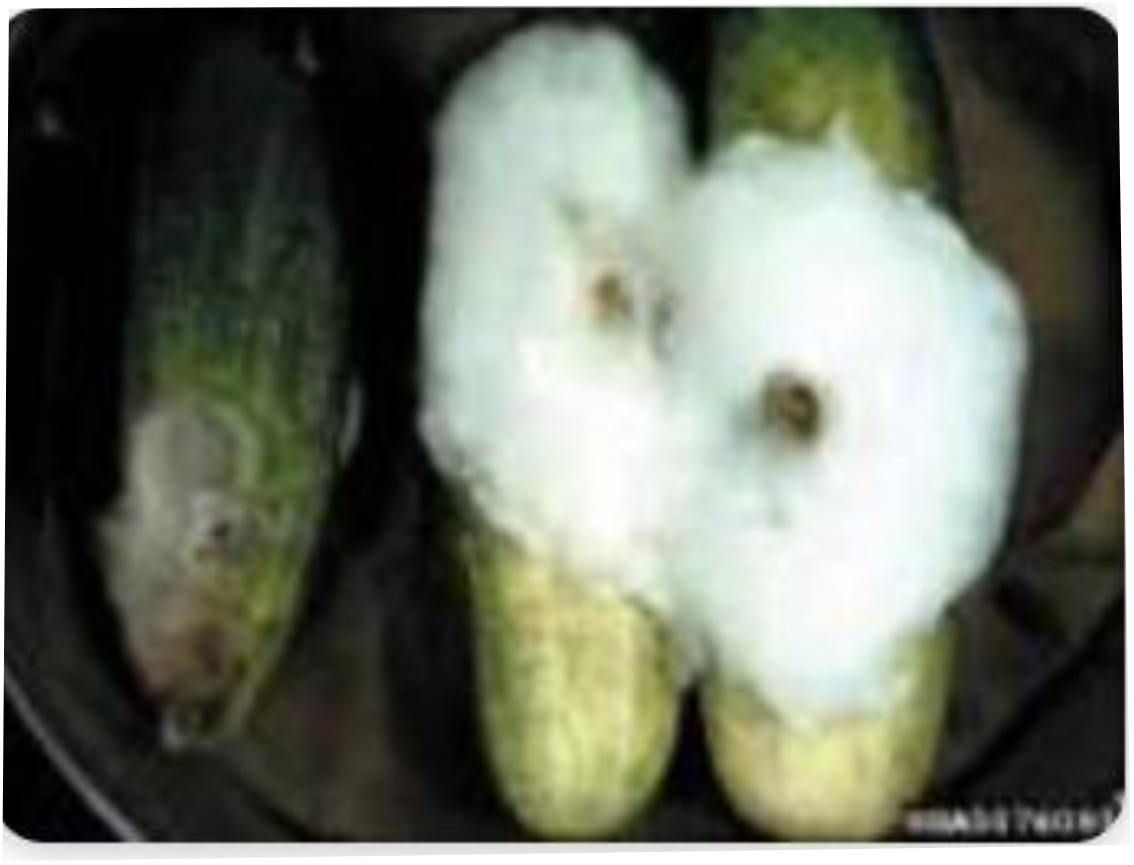
Rotten Cucumber fruit

**Fig. 2.**
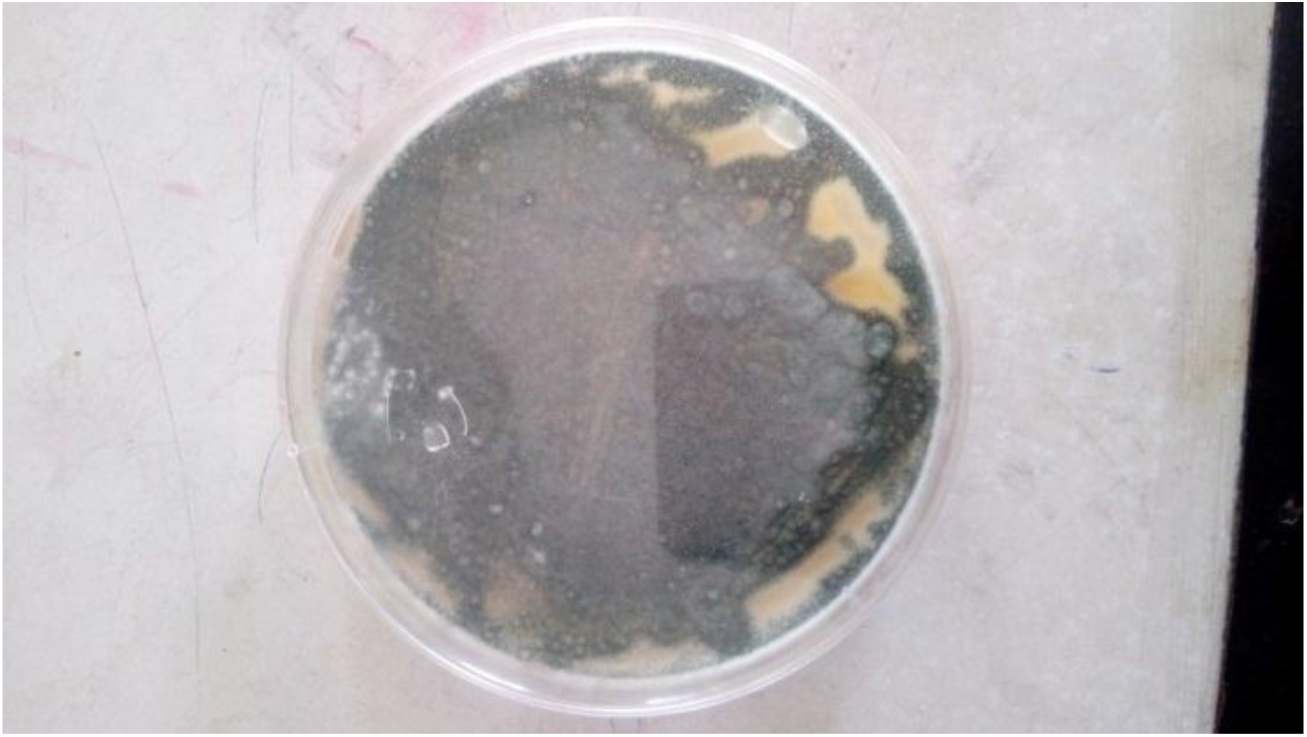
Pure culture plate *Penicillum* sp.

**Fig. 3.**
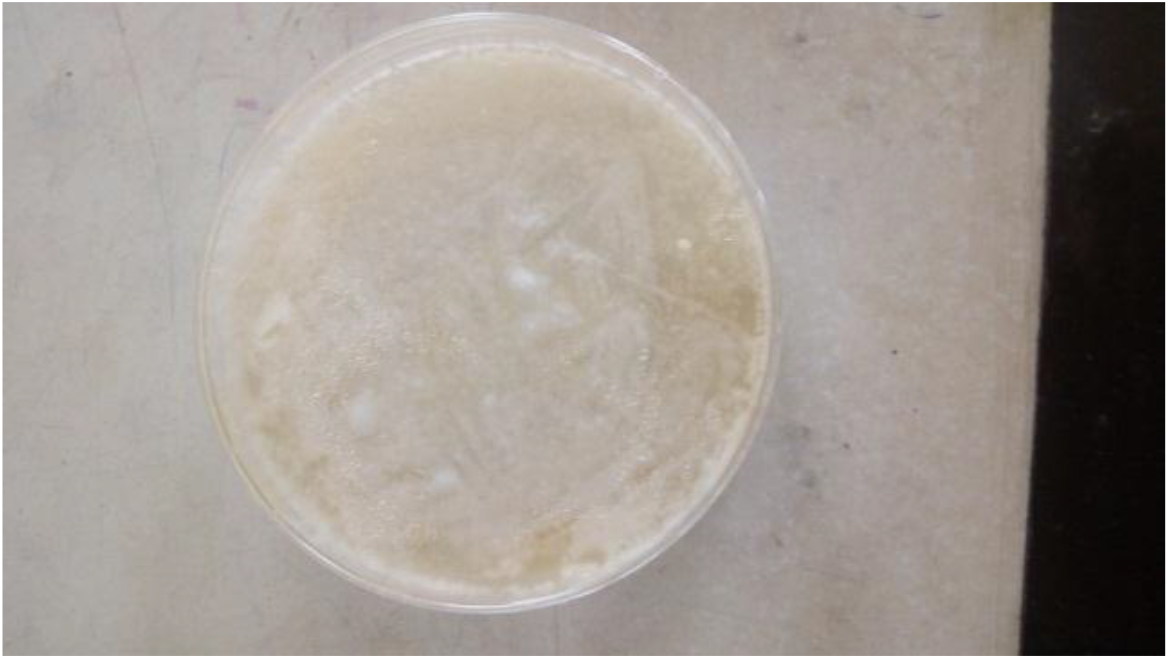
Pure culture plate *Fusarium oxysporium*

**Fig 4.**
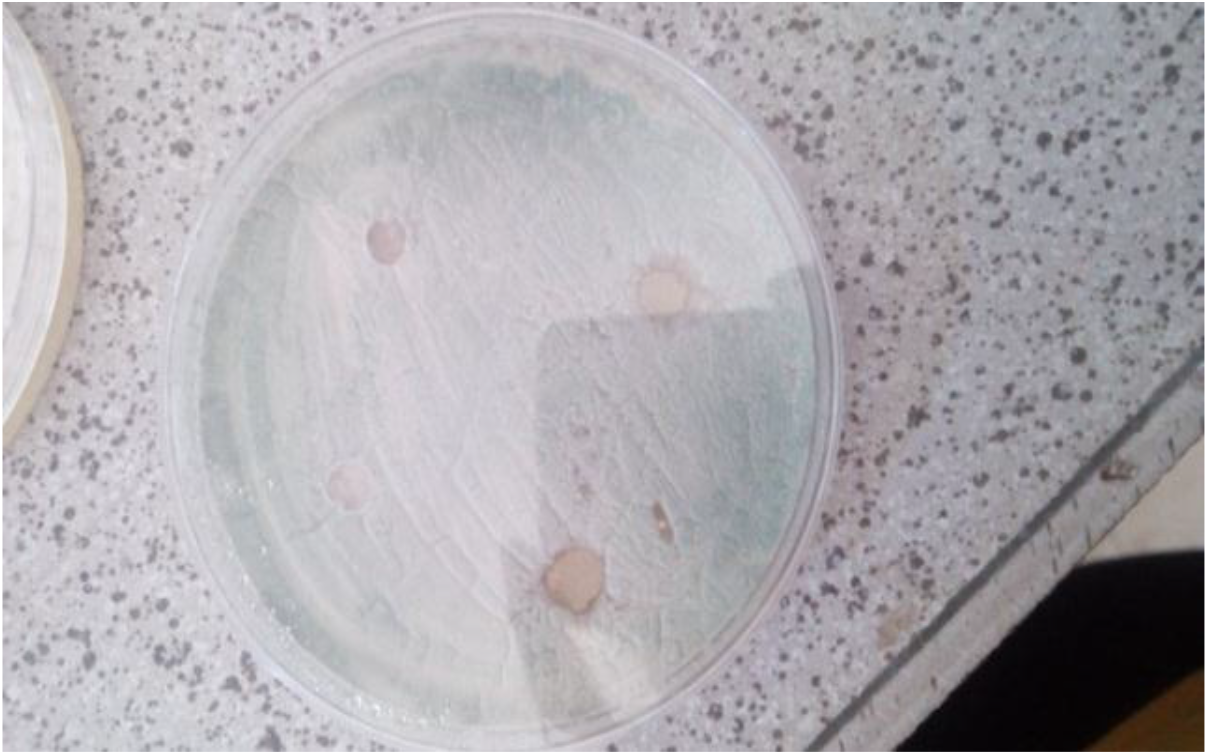
Lowest inhibitory effect on *Penicillum* sp.

**Fig 5.**
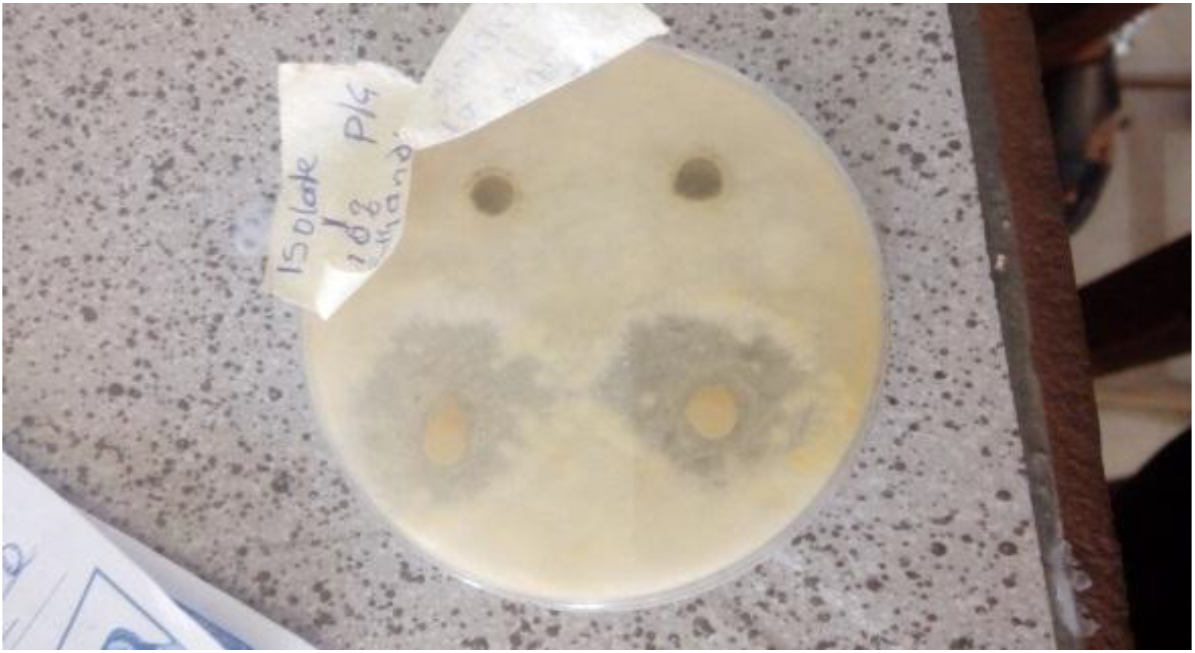
Highest Inhibitory effect on *Fusarium oxysporium*

**Fig 6.**
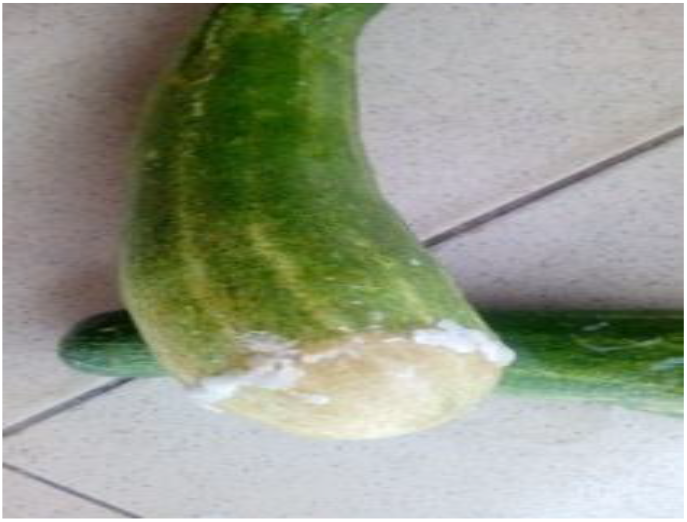
Pathogenicity result of *Penicillum* sp

**Fig 7.**
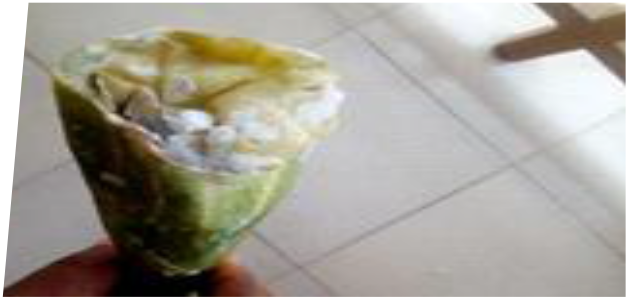
Pathogenicity result of *Fusarium oxysporium*

**Table. 1.**
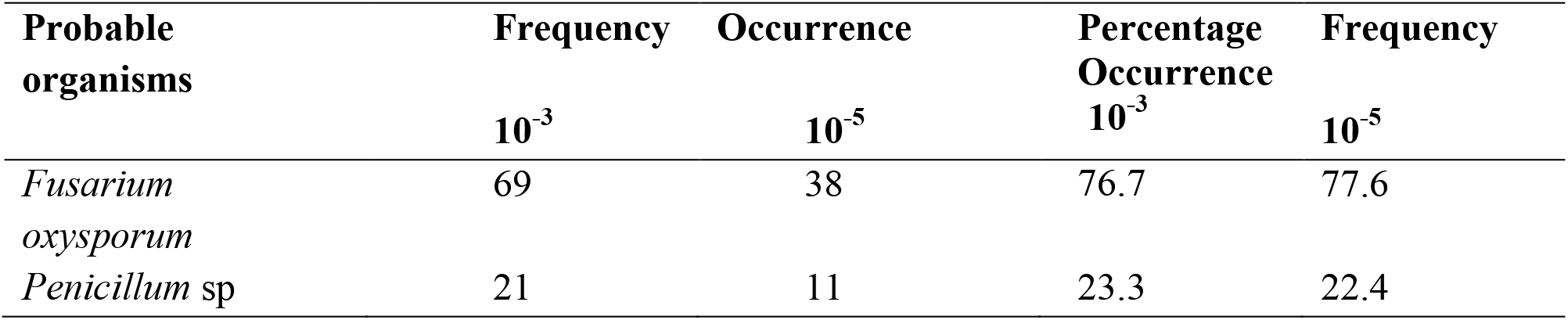
Total percentage frequency occurrence of the fungi isolated from rotted cucumber fruit.

**Table 2.** Zone of inhibition of plant extracts on the radical growth of *Fusarium oxysporium* and *Penncillum* sp.

| Isolates | Extracts | Mean zones of inhibition (mm) |  |  |
| --- | --- | --- | --- | --- |
|  |  | Concentration (%) |  |  |
|  |  | 70 | 100 | Control |
| *Fusarium oxysporium* | *Piper guineense* |  |  |  |
|  | Ethanollic extract | 17.0 | 26.5 | 0.0 |
|  | Aqueous extract | 0.0 | 8.5 | 0.0 |
|  | *Aframomum melegueta* |  |  |  |
|  | Ethanollic extract | 6.5 | 7.5 | 0.0 |
|  | Aqueous extract | 0.0 | 0.0 | 0.0 |
| Penicillum sp. | *Piper guineense* |  |  |  |
|  | Ethanollic extract | 12.5 | 18.0 | 0.0 |
|  | Aqueous extract | 4.0 | 12.0 | 0.0 |
|  | *Aframomum melegueta* |  |  |  |
|  | Ethanollic extract | 11.5 | 15.5 | 0.0 |
|  | Aqueous extract | 0.0 | 15.5 | 0.0 |

## 4.0 DISCUSSION

Different fungi were isolated from rotten cucumber fruit in accordance with those reported by Effiuvwevwere [14] who reported six (6) genera isolated from cucumber seed including *Fusarium oxysporum* and *Penicillium* sp. All these fungi reduced seed variation, *Fusarium oxysporum* caused a highly reduction in seed germination. The result showed variation of different fungal species and *Fusarium oxysporum* had the highest proportion of the fungal appearance in the rotten cucumber fruit where it recorded the highest percentage of frequency of 76.7% and 77.6% and the lowest appearance in the sample of cucumber was 23.3% and 22.4% which was recorded for *Penicillium* sp. The result is in line with the work of Effiuvwevwere [14] who reported *Fusarium oxysporum* as the most frequent occurring fungi found in rotten cucumber.

The result of the phytochemical screening of *Piper guineense* extracts shows the presence of saponins, flavonoids, alkaloids and cardiac glycosides, both in aqueous and ethanolic extracts and tested negative in tannins (Table 3). The phytochemicals of this study is consistent with the report of Nwankwo *et al* [15] who reported the most abundant phytochemicals in flavonoids and alkaloids and tannins tested negative in *Piper guineense* extracts. Biological function of flavonoids as one of the vital screened phytochemicals with high value in this study has protection against allergis and inflammation, free radicals, platelet aggregation, microbes [16].

**Table 3.**
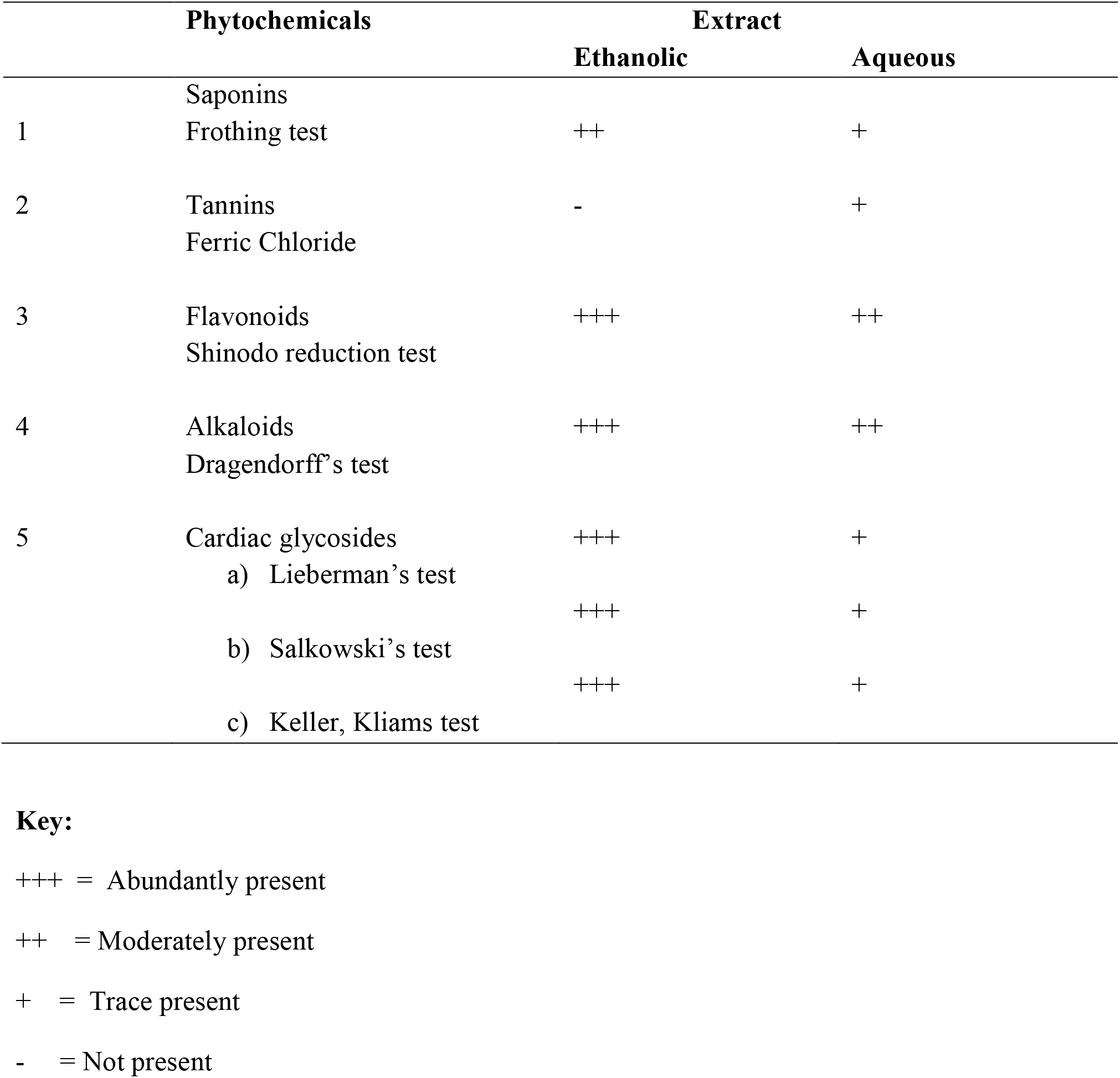
Phytochemical Screening of 70% ethanolic and aqueous extracts of *Piper guineense* (Ashanti pepper)

The antifungal activity of *Piper guineense* extracts under study showed varying zones of inhibition to the test fungal isolates. The ethanolic extract was more consistent than the aqueous extract in this study. However the highest zone of inhibitions was the ethanolic extract of *Piper guineense* extracts at 100% concentration on both *Fusarium oxysporum* and *Penicillium* sp. Which was 26.5mm and 18mm respectively (Table 3) while in aqueous extract both isolates recorded 8.5mm and 12mm. This finding with the comparative aqueous and ethanolic seed extracts of *Piper guineense* does not correlate with the studies of Nwinyi *et al* [17] where they reported that ethanolic extract of *Piper guineense* has less antifungal activity than the aqueous extract. This may be due to the solubility of the active compounds in water or the presence of inhibitors of antifungal components in ethanol. In this study, the aqueous extract of *Piper guineense* generally showed less antifungal activity than ethanolic extract against the isolates. The less activity of aqueous extract than ethanolic extract against most microbial strains investigated in this study is in agreement with previous work which shows that aqueous extract of plant generally show little or no antifungal activities [18]. It has been reported by Okigbo and Igwe [18] that inactivity of plant extracts maybe due to age of plant, extracting solvent, method of extraction and time of harvesting of plant materials. There are variations in the degree of antimicrobial activities of the extract on the isolates. The variation is due to difference in responses by the isolates to different active compounds present in the plant. Ethanol extracts of *Piper guineense* showed more antifungal activity against *Fusarium oxysporum* than *Penicillium* sp. The antifungal effect of the seed solvent extracts of *Piper guineense* may be attributed to the phytochemicals present in it. Distilled water which served as control showed no activity against the test organisms.

The results of the phytochemical screening of *Aframomum melegueta* (Alligator pepper) as presented in table 4 shows the presence of saponin, flavonoid, alkaloid cardiac glycoside in the ethanolic and aqueous extracts and tested negative in the tannins. The result is consistent with the report of Nwankwo *et al* [15] who reported the most abundant phytochemicals is alkaloid in *Aframomum melegueta* and tannins tested negative in both ethanolic and aqueous extracts.

**Table 4.** Phytochemical Screening of 70% ethanolic and aqueous extracts of *Aframomum melegueta*.

| Phytochemicals |  | Extract |  |
| --- | --- | --- | --- |
|  |  | Ethanolic | Aqueous |
| 1 | Saponins<br>Frothing test | ++ | + |
| 2 | Tannins<br>Ferric Chloride | - | - |
| 3 | Flavonoids<br>Shinodo reduction test | ++ | - |
| 4 | Alkaloids<br>Dragendorff's test | ++ | + |
| 5 | Cardiac glycosides |  |  |
|  | a) Lieberman's test | +++ | + |
|  | b) Salkowski's test | ++ | + |
|  | c) Keller, Kliams test | +++ | + |

+++ = Abundantly present
++ = Moderately present
+ = Trace present
- = Not present

The antifungal activity of *Aframomum melegueta* seed extracts was investigated against *Fusarium oxysporum* and *Penicillium* sp. isolated from rotten cucumber, using wet disc diffusion method. The result showed that the ethanolic extracts exhibited higher antifungal activity more than aqueous. The ethanolic extract at 70% and 100% concentration on *Fusarium oxysporum* showed inhibition zones of 6.5mm and 7.5mm respectively and at 70% and 100% concentration *Penicillium* sp showed 11.5mm and 15.5mm respectively (Table 4). This result showed that the ethanolic extract exhibited higher antifungal activity on *Penicillium* sp. Having the highest zone of inhibition (15.5mm and 11.5mm) than with *Fusarium oxysporum*. This observation is consistent with the report of Alo *et al* [19] who showed that ethanolic extract of *Aframomum melegueta* inhibited the growth of all the fungi tested. This observation is in agreement with the report of Doherty *et al* [20], who reported effectiveness of the seed extract of *Aframomum melegueta* against the tested organisms and suggested that the plant have broad spectrum activity. Similar observation has been reported by Oyagade *et al* [21].

Aqueous extract showed no zone of inhibition on both isolates and distilled water which served as control showed no activity against the test organisms. This report shows that ethanol is a better solvent for the extraction of active substance from the test plant part. This antifungal effect could be attributed to the phytochemical constituents of the plant seed such as flavonoids, saponins, cardiac glycosides and alkaloids. It also shows that the seed extracts of *Aframomum melegueta* can be used in the treatment of infections associated with the organisms. The combination of the two plants extracts showed no synergistic effect against the fungal isolates.

## 5.0 CONCLUSION

Ethanolic extracts of *Piper guineense* and *Aframomum melegueta* inhibited the growth of fungal pathogens of cucumber fruit rot, *Fusarium oxysporium* and *Penicillium* sp . at 100% and 70% concentrations. The inhibition of the growth of fungi causing cucumber rot is an indication that the extracts were effective in the control of cucumber rot. It is obvious that plant extract is currently considered as a potential botanical tool to solving the problem of plant rots. Thus, the antifungal potential of the plants extracts used in this study makes them suitable as botanical fungicides. Therefore, efforts should be geared towards harnessing the antimicrobial potentials of the vast plant resources to prevent plants rots

## Acknowledgements

The article was funded by TETFund research grant, Nigeria

## Disclosure of conflict of interest

The authors have declared no conflict of interest

## Notes

### Competing Interest Statement

The authors have declared no competing interest.

